# Optimization of process parameters for melanin nanoparticles synthesised from *Pseudomonas stutzeri* (BTCZ 109) using OFAT method and its anticancer property evaluation

**DOI:** 10.64898/2026.07.07.736906

**Authors:** Dayana Mathew, Sarita G. Bhat

## Abstract

Melanins are biological macromolecule with immense functionality synthesised by a wide spectrum of living organism. It is mainly synthesised by the oxidative polymerization of indolic and phenolic compounds through several enzymatic process. It has wide spread application in agriculture, cosmetic and therapeutic industry due to its various properties including antioxidation ability, UV protection efficiency and anticancer activity. Because of this wide range of application in different sectors, large scale production and commercialization attains enormous consideration. The present study deals with the effect of 12 different process parameters on melanin production viz., production media, incubation time, inoculum concentration, pH, temperature, agitation, carbon source, phosphate and magnesium source, CuSO_4_.5H_2_O, sodium chloride and L-tyrosine on melanin production by *Pseudomonas stutzeri* strain BTCZ 109 obtained from Arabian sea sediments was evaluated. After optimizing the important process parameters, the bacteria showed about ∼ 4.65 fold increase in melanin production compared to unoptimized cultural conditions. The melanin optimized through this method was found to be nano sized. The Nano sized DOPA melanin in treating Skin cancer cell line SK ML28 which showed a dose-dependent activity with an IC_50_ value of 164 μ g/mL. All these results highlight the therapeutic efficiency of DOPA melanin Nano particle as promising bioactive molecule.

**Highlights:**

- Process parameter optimization for large scale production of Nano-melanin obtained from *Pseudomonas stutzeri* strain BTCZ 109 through one-factor-at-a-time method.
- Optimization of 12 different process parameters were carried out which led to an increase in yield of ∼ 4.65 fold compared to unoptimized culture conditions.
- Its efficiency in treating skin cancer cell line SK ML28 was also detected.

## 1. Introduction

Melanins are a group of biological macromolecules produced primarily by the oxidation of indolic and phenolic compounds attains immense functionality among a wide range of prokaryotic and eukaryotic niches (Surwase *et al*., 2013). The macromolecular structure and chemical composition of melanin provides it with several physicochemical properties that allow them to act as ultraviolet absorbers, cation exchangers, drug carriers, amorphous semiconductors, X-ray and γ-ray absorbers (Lagunas Muñ oz *et al*., 2006). The biosynthetic pathway of melanin and melanin itself can be considered as potential target for antimicrobial drug discovery (Kiran *et al*., 2017). In recent years, melanin and related molecules were widely studied in biomedical field such as bioimaging, photothermal therapy and drug delivery vehicle (Hou *et al*., 2019). They are also widely employed in many related fields as functional material due to their excellent biodegradability and biocompatibility (Huang *et al*., 2018). The nano sized structural distribution of this enigmatic biomolecule obtained from bacterial strain *Pseudomonas stutzeri* BTCZ 109 was also appraised in our previous study which leads to its applicability in biomedical field.

The wide spread functionality of this biomolecule in pharmaceutical, cosmetological, agricultural and therapeutic industry increases its value industrially and which leads to optimization of process parameters for their large-scale production and commercialization. The cultural and fermentation conditions optimization, especially physical and nutritional condition are two of the key important parameters in the development and designation of any successful fermentation process leading to development of most economical and practically feasible bioprocess. Here in this study, the nano sized melanin particle obtained from novel isolate of *Pseudomonas stutzeri* from marine environment was employed for univariant experimental design technique through one-factor-at-a-time method to determine the most suitable cultural condition for increased bioproduction of this biomolecule.

Present study shed light on to 12 process parameters which helps to increase the production of nano-melanin particles from this particular strain of *Pseudomonas stutzeri* BTCZ 109. The important parameters considered here for increase in melanin production are production media, incubation time, inoculum concentration, pH, temperature, agitation, carbon source, phosphate and magnesium source, trace elements, sodium chloride and L-tyrosine. All these parameters were critically studied and evaluated to determine the most influencing factors affecting melanin production in bacterial strain. Minimal media (YMM) (Yabuuchi and Ohyama, 1972) containing only three salts (KH_2_PO_4_, NaCl and MgSO_4_) and additional 2gm/l of L-tyrosine was selected for the entire study considering economic feasibility and for reducing tedious steps during purification (Kurian *et al*., 2014). The purified nano-melanin particles thus obtained was then evaluated in Skin cancer cell line SK ML28 using microculture tetrazolium assay (MTT).

## 2. Materials and method

### 2.1 Microorganism and culture condition

#### 2.1.1 Microorganism

*Pseudomonas stutzeri* BTCZ109 (Gen Bank accession Number: MW479152) isolated from sediment sample collected during Sagar Sampada cruise no.305 from Arabian sea (9 6’ N, 75 22’ E) at 96 meter depth during a previous study and stored in the lab [Kurian et al, 2014] using paraffin over lay method was purified and further screened for extracellular melanin production in L-tyrosine agar plates.

#### 2.1.2 Chemicals used

L-tyrosine, synthetic DOPA melanin from Sigma, USA, Luria Bertani broth, Nutrient broth, peptone yeast extract, tryptone soya broth, Zobel’ s marine broth, and L-tyrosine basal broth, sodium chloride (NaCl), magnesium sulfate (MgSO_4_), potassium dihydrogen phosphate (KH_2_PO_4_), NaOH, 1N KOH, FeCl_3_, methanol, absolute ethanol, acetone, acetonitrile, ethyl acetate, 1-butanol, chloroform, benzene, 2-propanol, petroleum ether, hydrogen peroxide and sodium hypochlorite solution, starch, glucose, lactose, sucrose, glycerol, PBS solution. MTT assay cell line used; SK ML 28 cell line (Human skin cancer) (National Centre for Cell Sciences (NCCS), Pune, Dulbecco’ s Modified Eagles Medium (DMEM-Himedia), Fetal Bovine Serum (FBS), antibiotic cocktail containing Penicillin, Streptomycin, Amphotericin B, FBS and DMSO or Glycerol.

### 2.2. Optimization of melanin production media using One-factor-at-a-time Method

In the present study effect of 12 different process parameters were considered viz., production media, incubation time, inoculum concentration, pH, temperature, agitation, carbon source, nitrogen source, phosphate and magnesium source, trace elements, sodium chloride and L-tyrosine.

#### 2.2.2. Effect of production media on extra cellular synthesis of Nano-melanin pigment for the strain BT CZ 109

The most favourable media for the extra cellular melanin production was determined by inoculating initially with 10% of bacterial inoculum having 2* 10^9^ CFU/mL for BT CZ109 in six different media. The important media considered during this process were Luria Bertani broth, Nutrient broth, peptone yeast extract, tryptone soya broth, Zobel’ s marine broth, and L-tyrosine basal broth in 50 mL volume in a 250 mL Erlenmeyer flask with an additional supplement of 2% L-tyrosine and 1.5% NaCl. pH was adjusted to 7 and autoclaved. Melanin production was monitored spectrophotometrically at 400nm using synthetic melanin (Sigma, USA) as standard, the liquid medium that presented the highest yield of pigment was selected for further experiments (Kurian *et al*., 2015). The bacterial culture grown 9 days in liquid media was taken for calculating the yield. After 9 days of incubation at a temperature of 37 ° C and 140rpm in orbital shaker (Orbitek, scigenics, India) about 1 mL of the media was taken out and centrifuged at 8, 000 rpm (5000x g) at 4 ° C for 10 min to remove the cell debris. The supernatant was then taken out for determine the Nano-melanin production OD_400_ using UV-visible spectrophotometer (Shimadzu UV-1800) (Sajjan *et al*., 2013; Yabuuchi *et al*., 1972; Turick *et al*., 2002).

#### 2.2.3. Effect of incubation time on the extra cellular synthesis of melanin

For determining the optimum incubation time on the production of melanin 250mL Erlenmeyer flask with 50 mL media inoculated with 10 % inoculum were kept for melanin production in orbital shaker (Orbitek) at 140rpm and 37 ° C for 9 days of incubation (216 h). The production media was then monitored at a regular interval of 12h by determining the biomass OD_600_ and melanin production OD_400_ using UV-visible spectrophotometer (Shimadzu UV-1800). About 1 mL of the media was taken out and centrifuged at 8, 000 rpm (5000x g) at 4 ° C for 10 min to remove the cell debris (Kurian *et al*., 2015). The supernatant was then taken out and the melanin production was determined at OD_400_ using UV-visible spectrophotometer (Shimadzu UV-1800). All the experiments were performed in triplicates.

#### 2.2.4. Effect of inoculum concentration

Optimization of inoculum concentration can be evaluated in a minimal media (YMM) (Yabuuchi and Ohyama, 1972) containing only three salts (KH_2_PO_4_, NaCl and MgSO_4_) in 50 mL volume in a 250 mL Erlenmeyer flask with an additional supplement of 2% L-tyrosine and 1.5% NaCl by varying the inoculum concentration percentage. Here in this experiment 2-10% inoculum was taken and inoculated into 50 mL production media in 250 mL Erlenmeyer flask and incubated in shaker at 37 ° C and 140 rpm. The inoculum concentration that yielded maximum concentration of melanin was then determined using UV spectrophotometer and taken for further optimization studies. All the experiments were performed in triplicates.

#### 2.2.5. Effect of pH on the extracellular production of melanin pigment

The initial pH of the selected suitable liquid media that had been obtained from previous experiment was adjusted from 5.0 to 9.0 using 1N NaOH and 1N HCl before being autoclaved. After inoculating the culture media with the inoculum concentration already optimized was then incubated in a rotary shaker at 37 ° C and 140rpm for 5 days in the case of BT CZ109 without interception. The pH value that yields maximum melanin production was then selected for further optimization experiment.

#### 2.2.6. Effect of Carbon source on the extracellular production of melanin pigment

The effect of different carbon sources on melanin production was performed by using 10% (v/w) of 5 different carbon sources like, starch, glucose, lactose, sucrose(w/v) and glycerol(v/v), in 1x PBS solution and pH was adjusted to 8 using 1N NaOH. These carbon sources were then autoclaved at 10 lbs pressure for 10 min before inoculated. The samples taken were analysed for melanin production and the experiments were conducted at pH 8, agitation 140 rpm (Orbitek, Scigenics, India) and incubation temperature of 37 ° C. Melanin production media preparation, inoculum media preparation, quantification of melanin, inoculum preparation and extraction and purification of melanin were done as described earlier.

#### 2.2.7. Effect of Potassium dihydrogen orthophosphate

Optimization of potassium dihydrogen orthophosphate for maximum production of melanin was determined by using differing concentrations of potassium dihydrogen orthophosphate ranging from 0.5-2.5 g/l in the media and kept in orbital shaker at 37 ° C and 140rpm for 5 days of incubation in the case of BT CZ 109. Cell biomass and melanin production was then analysed by using UV spectrometer.

#### 2.2.8. Effect of agitation

The effect of different agitation rates on melanin production was determined by using100 to 180 rpm (Orbitek, Scigenics, India) and incubation temperature of 37 ° C. The production media was analysed after 5 days of incubation to determine the amount of melanin production as well as the biomass at OD_400_ for melanin production and OD_600_ for biomass determination using UV-visible spectrophotometer (Shimadzu UV-1800). About 1 mL of the media was taken out and centrifuged at 8, 000 rpm (5000x g) at 4 ° C for 10 min to remove the cell debris (Kurian *et al*., 2015). The supernatant was used to determine the melanin production at OD_400_ using UV-visible spectrophotometer (Shimadzu UV-1800). All the experiments were performed in triplicates.

#### 2.2.9. Effect of temperature

The effect of different temperatures ranging from 30-50 ° C at a regular interval of 5 ° C was employed to obtain the maximum amount of melanin production at already determined rpm. The amount of melanin production and biomass was monitored using UV-visible spectrophotometer (Shimadzu UV-1800). All these experiments were performed in triplicates.

#### 2.2.10. Effect of L-tyrosine concentration

L-tyrosine acts as the sole source of carbon and nitrogen in melanin production in optimization media. L-tyrosine at varying concentrations ranging from 2, 4, 6, 8, 10 g/L was used to determine maximum amount of melanin production and biomass concentration.

#### 2.2.11. Effect of Salinity

To determine optimum concentration of salinity, 250 mL Erlenmeyer flask having 50 mL of L-tyrosine basal medium with differing concentration of NaCl ranging from 0-3.5% (0.0.5, 1, 1.5, 2, 2.5, 3, 3.5 %) was used at optimizes temperature and agitation. All the experiments were performed in triplicates.

#### 2.2.12. Effect of MgSO_4_.7H_2_O

The effect of MgSO_4_.7H_2_O on melanin production on this particular strain of *Pseudomonas stutzeri* was determined by using 100mM concentration stock at different concentration ranging from 0.05-0.4 mM (0.05, 0.1, 0.2, 0.3, 0.4) was used. All the results obtained in triplicates was analysed using Graph pad prism software version 8.0 in windows 10.

#### 2.2.13. Effect of CuSO_4_.5H_2_O

Optimum concentration of CuSO_4_.5H_2_O was determined by using stock solution of 10mM CuSO_4_.5H_2_O after filter sterilization using 0.2 μ m filter (HiMedia, India) and at differing concentrations of CuSO4.5H2O ranging from 0.05-0.4 (0.05, 0.1, 0.2, 0.3, 0.4 mM). All the experiments were performed in triplicates and the results were analysed using Graph pad prism software.

### 2.3. Time course study after media optimization

By using one-variable-at-a-time approach initial screening of all the physical and chemical parameters allowing the maximum melanin production was performed. L-tyrosine acted as the sole source of carbon and nitrogen. 250 mL Erlenmeyer flask were inoculated with 8% inoculum and incubated in a shaking incubator with a shaking speed of 140 rpm at 35 ° C for 5 days. Samples were collected every 12 h and assayed for growth as well as melanin production.

### 2.4. Statistical analysis

Statistical analysis was carried out by two way analysis of variance (ANOVA) using Graph pad Prism version 8.0 for Windows 10. Test was used to determine significant difference (P< 0.05) Data is presented as means ± standard deviation. Differences were scored as statistically significant where *p* ≤ 0.05. For histogram and graphs, error bars represent standard deviations * P< 0.05, * * * P< 0.001, * * * * P< 0.0001. ns (non-significant).

### 2.5. Extraction and purification of Nano-melanin particle

The bacterial culture in the modified tyrosine-based medium was eventually de-cellularized by centrifugation after optimized period of melanin production [Narayanan *et al*., 2020]. The obtained melanin supernatant after centrifugation at 10, 000 RPM for 15 min. was acid precipitated using 0.1 N HCl. Further, the purity of melanin thus obtained was checked by dissolving in 1N NaOH using silica gel TLC (Thin layer chromatography) plates(Merck brand, Kenilworth, New Jersey) and a chromatogram was developed using the solvent system of n-butanol: acetic acid: water(70:20:10). The developed plates were then dried in an oven and the spots on the chromatogram were visualized with ninhydrin. The melanin pigment was then lyophilized and stored at − 20 ° C for future analysis (Keles *et al*., 2018)

### 2.6. Physical and Chemical properties of melanin granules

The physical and chemical properties of Bacterial melanin from BTCZ 109 was estimated compared to different solvents by keeping Synthetic DOPA-melanin as standard. Solubility test was performed against distilled water, 1M NaCl, 1N NaOH, 1N KOH, methanol, absolute ethanol, acetone, acetonitrile, ethyl acetate, 1-butanol, chloroform, benzene, 2-propanol and petroleum ether. Precipitation in 1N HCl and 1% (w/v) FeCl_3_ were determined. Reaction with oxidizing agents like 30% hydrogen peroxide and 10% sodium hypochlorite solution were determined (Suwannarach *et al*., 2019).

### 2.7 Anticancer property evaluation of melanin nanoparticle using MTT assay

Both safety and the anticancer activities of the purified melanin pigment of *Pseudomonas stutzeri* strain BTCZ109 were measured *in vitro* on skin cancer cell line SK ML 28. SK ML 28 cell line was used to determine the inhibitory effects of melanin pigment on cell growth using standard 3-(4, 5 dimethythiazol-2-yl)-2, 5-diphenyl tetrazolium bromide (MTT) assay. This colorimetric assay is based on the conversion of the yellow tetrazolium bromide (MTT) to a purple formazan derivative by mitochondrial succinate dehydrogenase in viable cells. The cells were cultured in Dulbecco’ s Modified Eagles Medium (DMEM-Himedia), supplemented with 10% heat inactivated Fetal Bovine Serum (FBS) and 1% antibiotic cocktail containing Penicillin (100U/ml), Streptomycin (100µ g/ml), and Amphotericin B (2.5µ g/ml). The cell containing TC flasks (25cm^2^) were incubated at 37° C at 5% CO_2_ environment with humidity in a cell culture incubator (Galaxy^®^ 170 Eppendorf, Germany). After incubation, the cells were treated with different concentration of melanin pigment (25, 50, 100, 150, 200 μ g/ml) and incubated for 24 h. After 24 h of drug treatment, 20 μ l of MTT solution at 5 mg/ml was added and incubated for 4 h. 100 μ l of Dimethyl sulfoxide (DMSO) in volume is added into each well to dissolve the purple formazan formed. The colorimetric assay is measured and recorded at absorbance of 570 nm using Eliza plate reader.

The relative cell viability in percentage was calculated and recorded as:

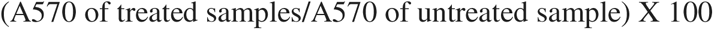

## 3. Results and discussion

### 3.1. Optimization of production media and the most influencing factors for its extracellular production

The media optimization was done with the help of One-factor-at-a-time method and the effect of different variables like media composition, incubation time, inoculum concentration, pH of the medium, Carbon source, KH_2_PO_4_ on growth and melanin production was determined.

#### 3.2. Effect of different media on melanin production

For determining the most suitable median for melanin production, the bacterial strain BT CZ109 was grown on different selected media like Luria Bertani broth, Nutrient broth, Peptone yeast extract, Tryptone soya broth, Zobel’ s marine broth and Yabuuchi and Ohyama minimal media with additional L-tyrosine and NaCl (Fig 1(a)). From the selected media the strain BT CZ 109 showed highest significant melanin production in Yabuuchi and Ohyama minimal media with additional L-tyrosine with a concentration of 1177.76+ /− 0.39 μ g/mL followed by ZMB (630.7+ /− 0.39 μ g/mL) and TSB (531.09+ /− 0.59 μ g/mL). The least melanin production was found in NB medium (221.129+ /− 0.71 μ g/mL).

**Figure 1.**
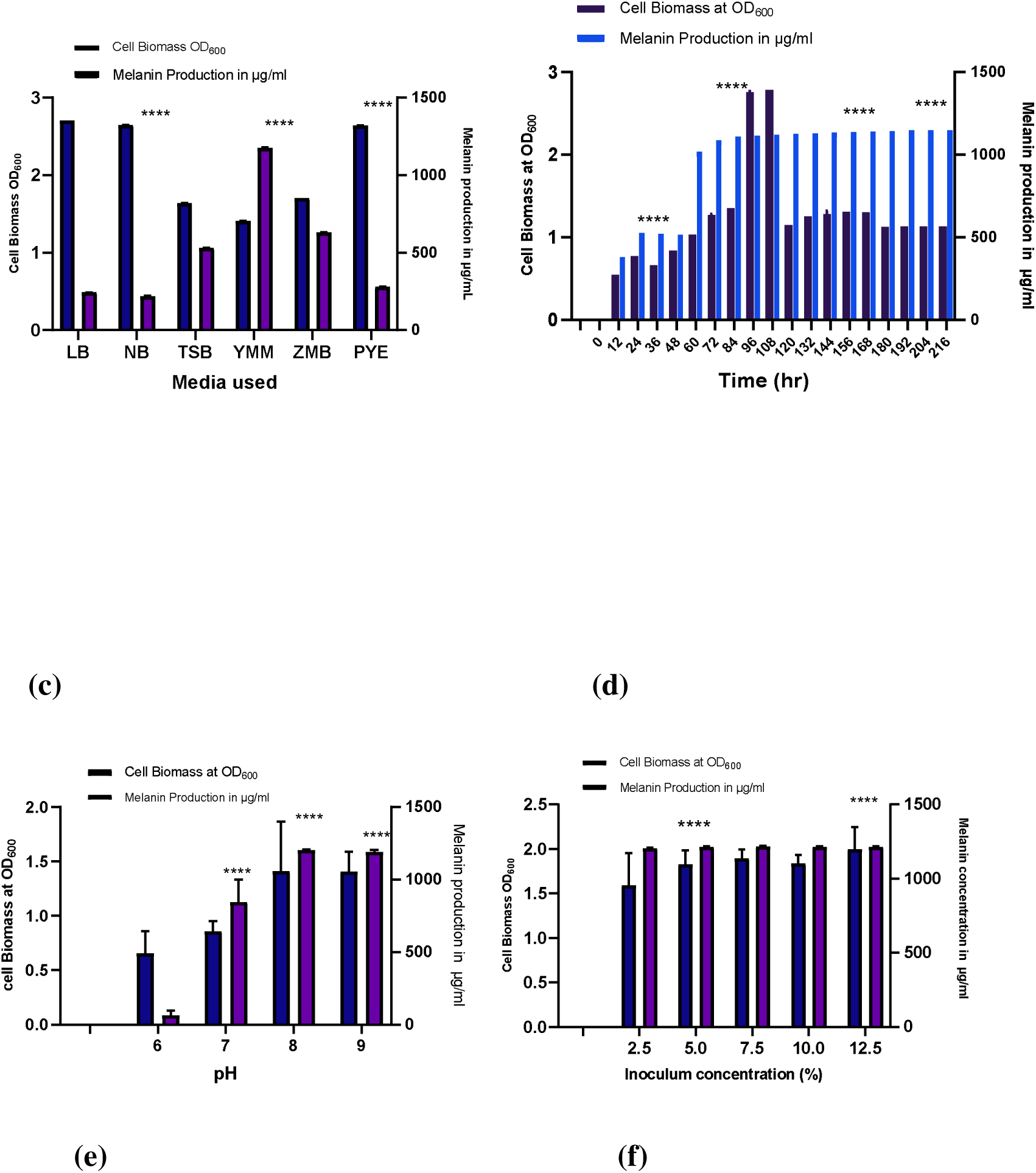
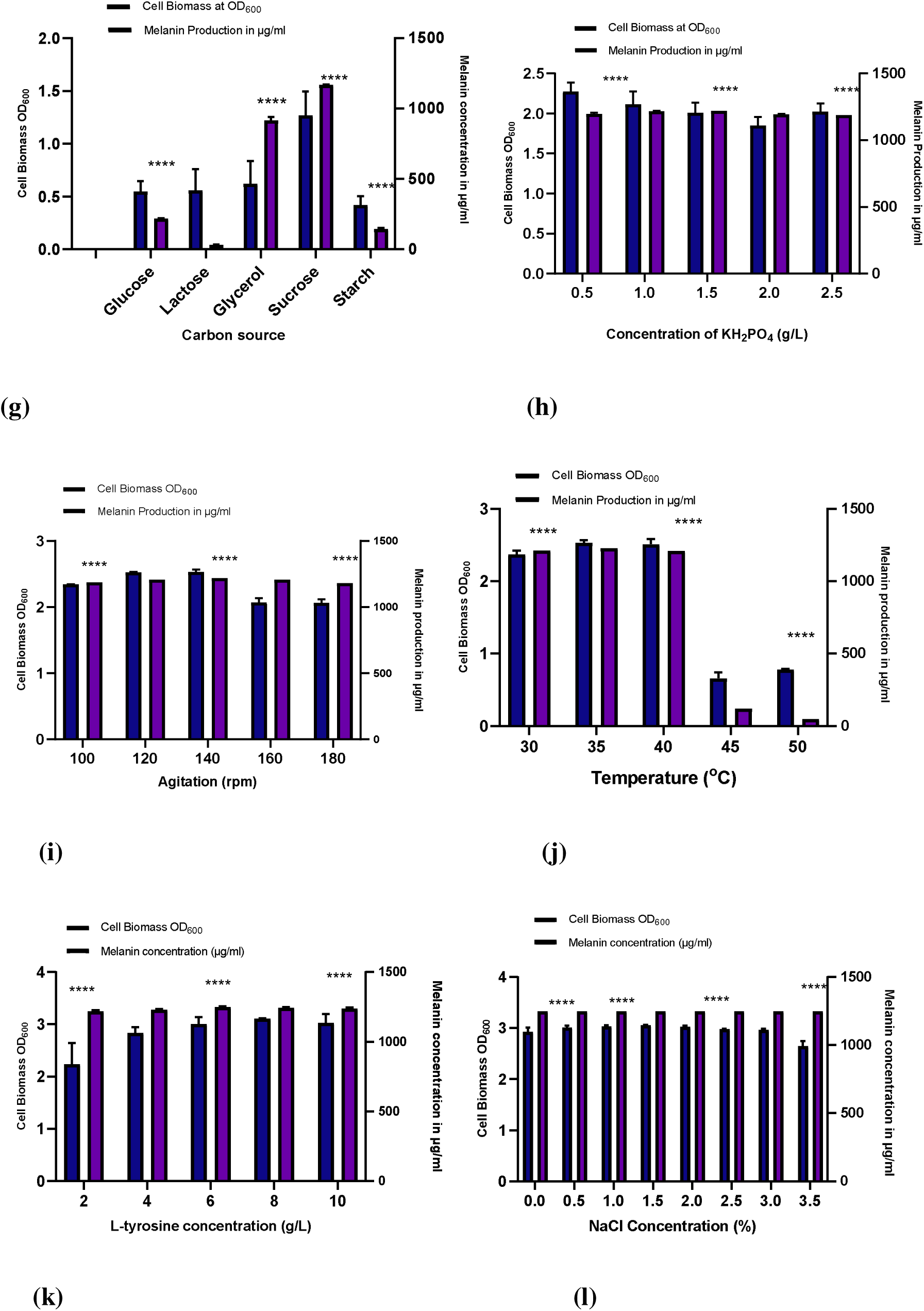
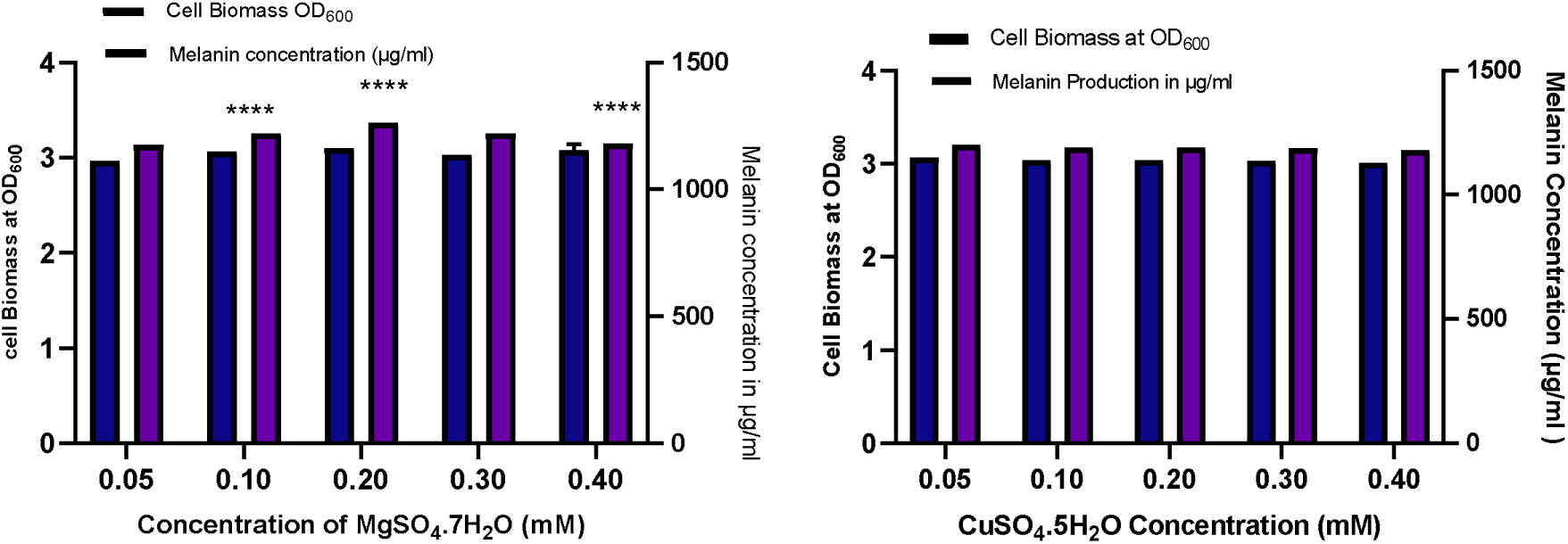
**(a)** Effect of different media on melanin production and cell biomass of bacterial strain BT CZ109 (b) Incubation time (c) pH (d) inoculum concentration (e) carbon source (f) KH_2_PO_4_ (g) temperature (h) L-tyrosine (i) NaCl (j) MgSO_4_.7H_2_O (k) CuSO_4_.5H_2_O. All the values were statistically analysed using Graph pad prism 8 in Windows 10. The results are means of three replicates +-SD For histogram and graphs, error bars represent standard deviations * P< 0.05, * * * P< 0.001, * * * * P< 0.0001. ns (non-significant).

Among these YMM was used further because of the less complexity of the medium and less tedious processes during purification. This is followed by ZMB with a production concentration of 1135.54+ /− 10.22 μ g/mL and TSB (1021.67+ /− 0.577 μ g/mL). The least significant melanin production was observed in NB (1002.077+ /− 38.62 μ g/mL).

#### 3.3. Effect of incubation time on melanin production

The highest significant melanin pigment yield was determined by taking sample and monitoring the melanin production using UV-Vis spectrometer at a regular interval of 12 h (Fig 1(b)). Notably, after an incubation time of 216 h the bacterium BTCZ109 showed maximum melanin production of 1150+ /− 30 μ g/mL respectively, and it was observed from the result that bacterial strain exhibited maximum melanin production in its stationary phase of growth.

#### 3.4. Effect of pH on melanin production

In the present study, both these microbial strains showed maximum growth and melanin production in alkaline pH range Fig 1(c). For the bacterial strain BTCZ109 the maximum melanin production attained at pH 8 was 1204.553+ ___ 2.992 μ g/mL. When reaching the acidic range the melanin production was found to be diminished. The maximum amount of melanin produced by this particular microorganism was about 1202.076+ ___ 5.65 μ g/mL.

#### 3.5. Effect of inoculum concentration on melanin production

The effect of different percentage of inoculum contributing to maximum melanin production and biomass concentration was detected against BTCZ109 (2* 10^9^ CFU/mL) fig 1(d). From the results it was inferred that the strain BTCZ109 showed maximum melanin production at 8% inoculum (1218.28+ /− 1.92 μ g/mL) and the minimum melanin production was shown by inoculum size of 2% (1204.95+ /− 4.32 μ g/mL).

#### 3.6. Effect of Carbon source

Different carbon sources like glucose, lactose, starch, sucrose, starch(w/v) and glycerol(v/v) were employed for determining the most effective carbon source for maximum growth and melanin production in the media (fig 1(e)). From the results it was inferred that bacterial strains BTCZ109 showed maximum growth and melanin production in media with Sucrose as carbon source. The concentration of melanin produced by BTCZ109 was about 1167.693+ /− 2.66 μ g/mL. But any of these results show remarkable increment in melanin production compared to the previous optimization step. So, from the result it was concluded that even the sugar, sucrose which showed maximum production cannot be considered as good candidate during optimization study.

#### 3.7. Effect of potassium dihydrogen orthophosphate

Melanin production and cell biomass concentration was monitored at varying concentration of potassium dihydrogen orthophosphate (fig1(f)) to determine the concentration at which melanin production reaches at its maximum. But from the result it was observed that potassium dihydrogen orthophosphate doesn’ t influence significantly on melanin production.

#### 3.8. Effect of agitation

The effect of aeration on melanin production was quantified using different agitation rates from 100-180 rpm (fig 1(g)) and from the results it was inferred that at 140 rpm the organism showed maximum melanin production along with previously optimized parameters and the concentration of melanin produced was about 1220.5+ /− 3.96 μ g/mL. Aeration is a significant factor which contribute in maximum melanin production

#### 3.9. Effect of temperature

Fig. 1(h) shows the effect of temperature on melanin production and biomass trend of *Pseudomonas stutzeri* strain. From the result it was observed that the particular strain is mesophilic in nature and amount of melanin production reached its optimum value at 35 ° C. Increase in temperature above 40 ° C causes considerable decrease in melanin production.

#### 3.10. Effect of L-tyrosine concentration

Fig.1(i) indicates the influence of L-tyrosine on melanin production, and from the graph it was clear that tyrosine concentration of 6 g/L showed the maximum amount of melanin production in this particular strain of *Pseudomonas stutzeri* and it was found to be about 1249.26+ /− 4.5 μ g/mL. Rapid increase in biomass concentration can also be noticed while using high concentration of L-tyrosine which helps the cells to attain stationary phase in a reduced time interval.

#### 3.11. Effect of Salinity

Effect of salinity on melanin biosynthesis was determined (Fig.1(j)] by using a wide range of NaCl concentrations ranging from 0-3%. From the results it was inferred that NaCl does not influence considerably on melanin production by *Pseudomonas stutzeri* BTCZ 109 strain.

#### 3.12. Effect of MgSO_4_.7H_2_O

Optimum MgSO_4_.7H_2_O concentration influencing Melanin production was determined (Fig.1(k)) and from the result it was observed that 0.2mM concentration showed maximum amount of melanin production of about 1263.64+ /− 0.16 μ g/mL.

#### 3.13. Effect of CuSO_4_.5H_2_O

From Fig.1(l) it was evident that CuSO_4_.5H_2_O does not significantly contribute much on increase in melanin production. The main enzyme involved in melanin biosynthesis was found to be laccase in our previous study using Sodium azide as inhibitor.

### 3.2. Time course study after media optimisation

The time course was studied up to 120 h of incubation in optimized medium and maximum melanin production of 1270.5 μ g/mL was achieved in 96 h of incubation (Fig.2). Similar to the earlier reports (Manivasagan *et al*., 2013) it was observed that melanin production built up slowly during exponential phase and attained maxima at the onset of stationary phase. But in the case of Biomass concentration, we can observe that log phase diminished considerably after media optimization. In the present study, the melanin production increased steadily and reached the maximum at 96 h of incubation in this particular strain of bacteria.

**Fig 2.**
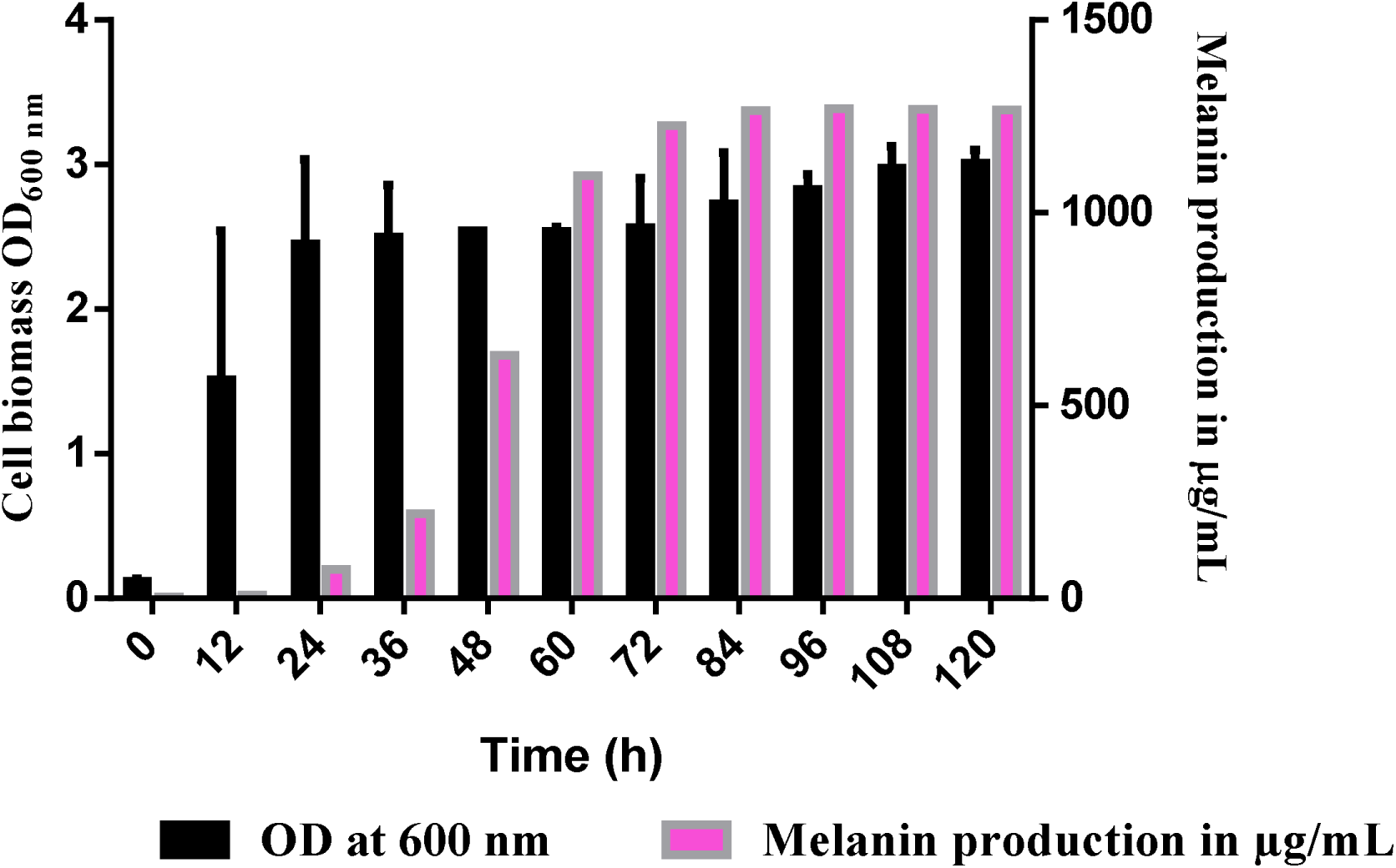
Time dependent production of melanin and biomass at a regular interval of 12 h was shown on the graph

### 3.3. Extraction and purification

Extraction of melanin nano particle was done by using 1 N HCl for 1 week of incubation and the purity check was performed by using Thin layer chromatogram using silica gel plates(Fig 3). The R_f_ value was found to be 0.6 cm which was similar to the R_f_ value of synthetic DOPA melanin.

**Fig 3.**
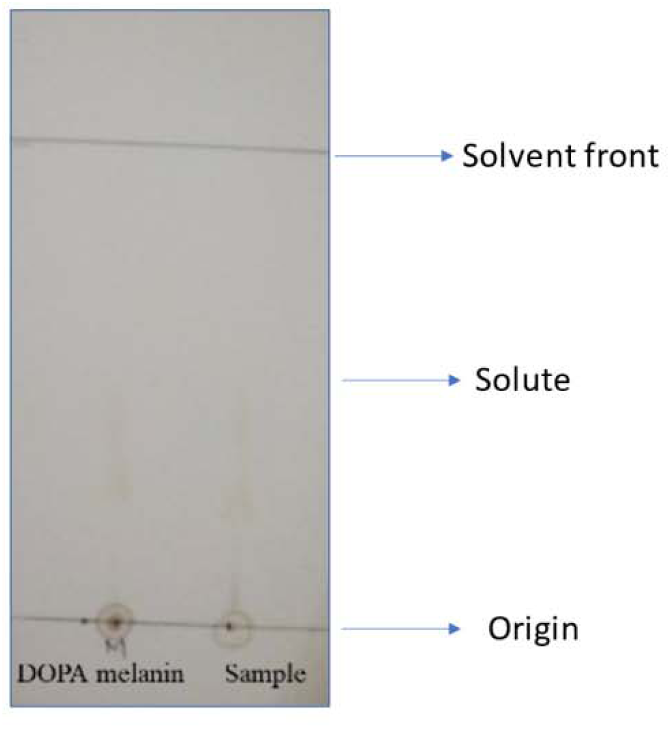
Silica gel plates showing chromatogram of synthetic DOPA melanin and bacterial nano-melanin

### 3.4. Physical and Chemical properties of melanin

The physical chemical properties of melanin obtained from marine bacteria BTCZ 109 was determined with the help of different solvents by keeping Synthetic DOPA melanin a standard (Table 1). From the physical and chemical property determination it was revealed that all these melanins were insoluble in distilled water, 1M NaCl, methanol, absolute ethanol, acetone, acetonitrile, ethyl acetate, 1-butanol, chloroform, benzene, 2-propanol and petroleum ether and soluble in 1N NaOH, 1N KOH. From the precipitation study conducted with 1N HCl and 1% (w/v) FeCl_3_, it was concluded that the pigment obtained from BTCZ109 get precipitated and showed similar result compared to Synthetic DOPA melanin standard. The oxidising property of the pigment showed positive result compared to synthetic melanin. This study conformed the similarity of bacterial melanin to synthetic DOPA melanin standard. The results obtained were similar to that of previous findings (Suwannarach *et al*., 2019) and it is tabulated as below.

**Table 1:**
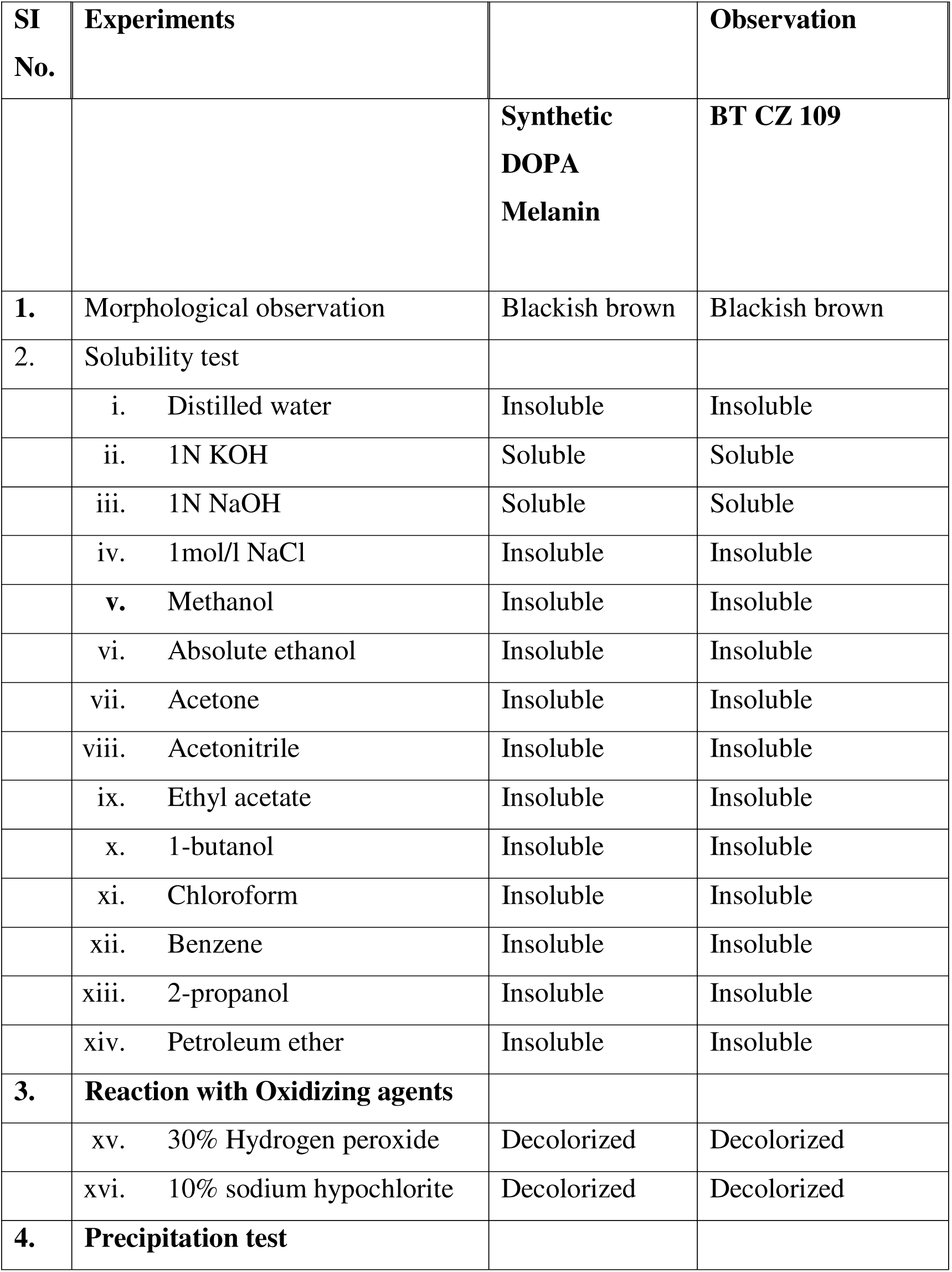
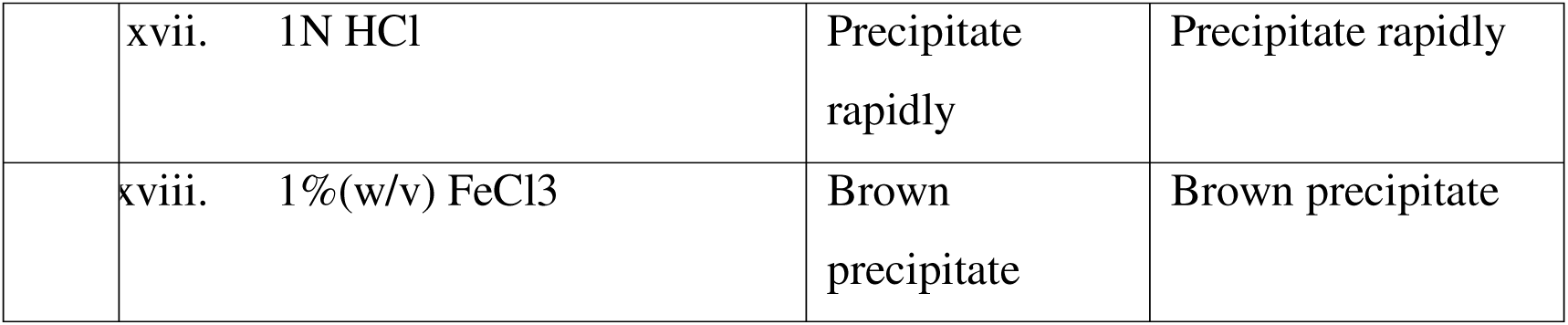
Physical and chemical properties of Bacterial Melanin (Melanin obtained from BT CZ109 and synthetic DOPA melanin)

### 3.5 Anticancer property evaluation of melanin nanoparticle using MTT assay

The safety pattern of the purified melanin pigment of *Pseudomonas stutzeri* strain BTCZ 109 was assayed on skin cancer cell line (SK ML28) using MTT assay. The obtained results were expressed as growth inhibitory concentration (IC_50_) values, which represent the melanin pigment concentration required to produce a 50% inhibition of cell growth after 24 h of incubation, compared to untreated control. A concentration dependant reduction in viability of cells was observed as compared with the untreated control, Cell shrinking and reduction in cell number can be observed at concentration range 150 and 200 μ g/ml in the fig 4(a). This indicates that the sample shows anticancer property at higher concentrations. The IC_50_ value was found to be 164 μ g/ml.

**Fig 4.**
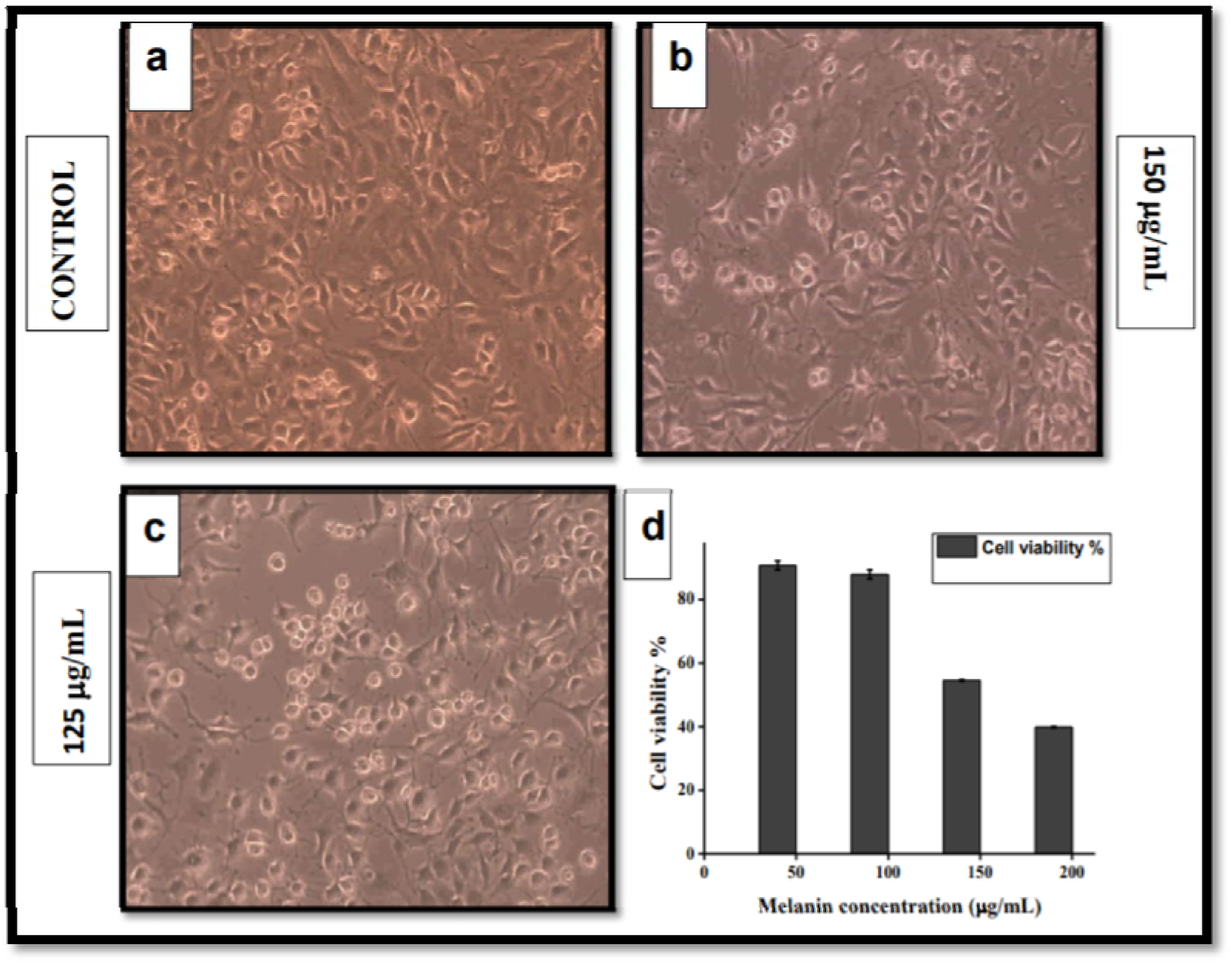
(a, b, c) Concentration dependendent reduction cell viability can be observed for control, 150 μ g/mL, 125 μ g/mL respectively (d) Graph shows the reduction in cell viability with increase in concentration of melanin.

## 4. Discussion

Melanins can be considered as natural biopolymers with several biomedical application including drug carrier, biosensor, molecular imaging agent etc. Here in the present study Nanosized melanin granules obtained from *Pseudomonas stutzeri* strain BTCZ 109 isolated from Arabian sea sediment sample was critically studied for evaluating its bioproduction, physical and chemical properties in several different solvents and finally its role in irradiating malignant melanoma. In microbes, especially in bacteria melanins were synthesised to tolerate against natural stress conditions ranging from ultraviolet radiation and toxic heavy metals to oxidative stress (Pavan *et al*., 2020). So, in our study, we tried to recreate similar stress by providing a minimal media with diminished nutrients. This nutrient depletion induced the bacteria to produce melanin at its stationary phase of growth. So far, several different kinds of bacteria including *Aeromonas salmonicida, Azotobacter, Mycobacterium, Micrococcus, Bacillus, Legionella, Streptomyces, Rhizobium, Vibrio, Proteus, Azospirillum, Pseudomonas aeruginosa, Hypomonas sp, Burkholderia cepacia, E. coli, Bordetella pertusis, Campylobacter jejuni, Yersinia pestis* etc were reported to be efficient melanin producers. Melanin production from marine bacteria was also a fascinating topic with limited literature. So far, bacteria like *Vibrio cholerae*, *Shewanella colwelliana* species capable to produce pyomelanin through tyrosine degradation pathway were reported. In another study on marine bacterium genus *Alteromonas* and *Marinomonas mediterranea* MMB-1T which belong to the phylum Proteobacteria as well as thermo-alkaliphilic *Streptomyces* isolated from limestone quarries of the Deccan traps were considered as excellent melanin producers (Tarangini *et al*., 2013).

Classical One-factor-at-a-time method was employed to scrutinize important factors and its level in melanin biosynthesis. 12 different process parameters including both physical and chemical factors were choosen for optimization study. From the selected factors some of them contributed significantly to melanin production but some were not. The factors like Incubation time, Inoculum concentration, pH, temperature, L-tyrosine, agitation and MgSO_4_.7H_2_O are factors which contribute significantly in increasing melanin production. On the other hand, KH_2_PO_4_, salinity, CuSO_4_.5H_2_O doesnot contribute much to production.

Considering the incubation time of melanin production, here we have taken 5days as optimum because after 86 h of incubation the cell biomass reaches at its stationary phase of growth and also the melanin production hits at its maximum after that. Similarly, a study conducted by El-Naggar *et al*., 2017, 6 days of incubation was considered to be optimum. Similar comparable results were observed in previous studies conducted by Vasanthabharathi *et al*., 2011.

In the present condition 2* 10^9^ CFU/mL in 5% inoculum for 5 days of incubation can provide 1270.5 μ g/mL of melanin. In a similar study conducted by (Sivaperumal *et al*., 2014), the inoculum concentration was fixed at 5% with a cell density of about 3.1* 10^4^– 5.4* 10^4^ CFU/mL. While performing pH optimization it was noticed that a slightly alkaline pH favours melanin formation in *Pseudomonas stutzeri* strain. One study conducted by Aghajanyan *et al*., 2005 suggested that pH range from 7-8 was required for *Bacillus thuringiensis* melanin synthesis. Melanin from marine Streptomyces sp. (MVCS13) showed maximum melanin production at a pH of 7.4 (Sivaperumal *et al*., 2014). Some previous research show evidence of bacterial melanin stable only in the pH range of 4.0-11.0 (Wang *et al*., 2000). Among different parameters studied pH of the growth medium plays a crucial role in the stability of melanin granule in the medium. pH of the medium helps in aggregation as well as disintegration of the granules. This keeps the granules in stable nano form after production in the media.

While considering the temperature optimization part of the study we can notice that the bacterium *Pseudomonas stutzeri* produces amount of melanin at 35 ° C which clearly defines that the bacteria is mesophilic aerobic non spore forming bacteria. In a similar study conducted on *Bacillus thuringiensis* (Ruan *et al*., 2004) the high melanin yield was obtained when temperature elevated to 42 ° C. In a recent study conducted on actinobacteria *Dietzia schimae* (Eskandari *et al*., 2020) maximum amount of melanin was obtained at an optimum temperature of 32 ° C. From all the results it was clear that temperature possessed significant role in melanin production in microorganisms especially bacteria. An agitation rate of 140 rpm is enough for providing sufficient aeration for the bacteria to proliferate and synthesis melanin.

L-tyrosine act as the primary precursor in melanisation in *Pseudomonas stutzeri* strain and also act as the sole source of carbon and nitrogen. From our previous study (not published yet) it was inferred that melanin formation in this bacterium was mainly occurs through laccase or polyketide synthases as well as homogentisic acid polymerization, yet little is known about this mechanism. The involvement of laccase in melanin synthesis was reported previously. While laccase is widely distributed in plants and fungi, its activity in melanin formation, lignolysis or detoxification, and related activity of this enzyme in bacteria has been rarely documented. Both laccase enzyme and tyrosinase possesses three conservative histidine residues and also possess similar function, but irrespective of that, it is activated under different conditions in different species. The enzymes other than laccase, like phenylalanine 4-monooxygenase, pterin-4-alpha-carbinolamine dehydratase, aromatic amino acid aminotransferase, and 4-hydroxyphenylpyruvate dioxygenase (Drewnowska *et al*., 2015) were also involved melanin formation in bacteria. But its participation requires further investigation.

MgSO_4_.7H_2_O at a very low concentration favours melanin formation by initiating DNA replication and cell division, whereas increase in concentration of MgSO_4_.7H_2_O above this value was detrimental for the organism. Surwase *et al*., 2014 reported slight increase in melanin production by *Bacillus sp*. with slight increase in MgSO_4_.7H_2_O.

Optimization of all these parameters finally leads to increase in melanin production ∼ 4.65 fold compared to unoptimized condition. The melanin so formed in the medium was nano sized and has got wide spread application therapeutically.

## 5. Conclusion

Melanins are a group of enigmatic biopolymers synthesised by all forms of life. Its therapeutic potential and cosmetological relevance make it into valuable bioproduct industrially. The nano sized structural distribution enhances its applicability in pharmaceutical field. Melanin can be used a potent drug carrier, molecular imaging agent as well as biosensors. Large scale production and commercialization attains immense importance at this scenario. Scaling up of the biomolecule through process optimization by considering both physical and chemical parameters attract more attention. Formulation of economically feasible media and selection of most relevant strain are challenging. Our study shed light on to process optimization through one-factor-at-a-time method on *Pseudomonas stutzeri* derived nano melanin and its application. Because of the possible applications, it can be considered as alternative to commercial pigment, thus, it is worth further investigation.

## 6. Acknowledgements

The first author acknowledge the support of Council of Scientific and Industrial Research (CSIR), Govt. of India for granting fellowships during the study (CSIR Award no: 09/239(0535)2018-EMR-1).

## 8. Author contribution

DM did all the experimental work and wrote the draft of the manuscript under the supervision SGB, who is her doctoral supervisor, and mentor, who modified the manuscript for publication.

